# Quantitative T_1_-relaxation corrected metabolite mapping of 12 metabolites in the human brain at 9.4 T

**DOI:** 10.1101/2021.12.24.474092

**Authors:** Andrew Martin Wright, Saipavitra Murali-Manohar, Anke Henning

**Affiliations:** High-Field Magnetic Resonance, Max Planck Institute for Biological Cybernetics, Tübingen, Germany; IMPRS for Cognitive & Systems Neuroscience, Tübingen, Germany; F.M. Kirby Research Center for Functional Brain Imaging, Kennedy Krieger Institute, Baltimore, Maryland, USA; Advanced Imaging Research Center, UT Southwestern Medical Center, Dallas, Texas, USA

**Keywords:** Quantitative, MRSI, metabolite, mapping, UHF, ultra-high field, spectroscopic imaging, spectroscopy

## Abstract

Magnetic resonance spectroscopic imaging (MRSI) is a non-invasive imaging modality that enables observation of metabolites. Applications of MRSI for neuroimaging applications has shown promise for monitoring and detecting various diseases. This study builds off previously developed techniques of short TR, ^1^H FID MRSI by correcting for T_1_-weighting of the metabolites and utilizing an internal water reference to produce quantitative (mmol kg^−1^) metabolite maps. This work reports and shows quantitative metabolite maps for 12 metabolites for a single slice. Voxel-specific T_1_-corrections for water are common in MRSI studies; however, most studies use either averaged T_1_-relaxation times to correct for T_1_-weighting of metabolites or omit this correction step entirely. This work employs the use of voxel-specific T_1_-corrections for metabolites in addition to water. Utilizing averaged T_1_-relaxation times for metabolites can bias metabolite maps for metabolites that have strong differences between T_1_-relaxation for GM and WM (i.e. Glu). This work systematically compares quantitative metabolite maps to single voxel quantitative results and qualitatively compares metabolite maps to previous works.

**Highlights:**

- Quantitative metabolite maps of 12 metabolites human brains acquired at 9.4 T.
- Voxel-specific T_1_-weighting corrections for water and metabolites.
- Comparisons of T_1_-weighted and T_1_-corrected metabolite maps.

## 1. Introduction

Proton magnetic resonance spectroscopic imaging (^1^H MRSI) is a non-invasive imaging modality that allows for direct observation of the distribution of metabolites. MRSI merges single voxel spectroscopy (SVS) with traditional MRI, and can be used to visualize the distribution of metabolites in given tissue types. Similarly to SVS studies, higher field strengths improve the detection capabilities of MRSI due to increased SNR and improved spectral resolution or by reducing acquisition times (Bogner et al., 2012; Henning et al., 2009; Henning, 2017). In regards to diagnostic imaging, MRSI aids in detection and monitoring of multiple neurodegenerative diseases such as multiple sclerosis (Hangel et al., 2018), glioma (Gruber et al., 2017), epilepsy (Pan et al., 2013), and more (Maudsley et al., 2020; Öz et al., 2014).

To date, quantitative ^1^H MRSI results have been shown with limited metabolite coverage and limited transparency in metabolite map quality. Most early works reported quantitative results in mM quantities for N-acetyl aspartate (NAA), tCr, and tCho (Bonekamp et al., 2010; Degaonkar et al., 2005; Hetherington et al., 1996; Horská et al., 2002; Pan et al., 1998; Soher et al., 1996). Three more studies included mI (Lecocq et al., 2014; Tal et al., 2012); one of which also included Glx (Mclean et al., 2000). The most recent quantitative work that expanded the metabolite coverage significantly was (Hangel et al., 2021) and reported quantitative results for 12 metabolites and showed metabolite maps for NAA, tCr, tCho, Glu, and mI.

Many studies targeting the human brain have been done at ultra-high field (UHF, ≥ 7 T) and have shown that higher field strengths improve MRSI data quality (Gruber et al., 2017; Henning, 2017; Nassirpour et al., 2016; Trattnig et al., 2016), and highlighted the benefit of ultra-short echo time (TE) sequences (Bogner et al., 2020; Kirchner et al., 2017; Nassirpour et al., 2016; Považan et al., 2015). Free induction decay (FID) MRSI benefits acquisitions by reducing the amount of signal loss due to T_2_-relaxation as well as reducing the amount of effects arising from J-coupling interactions. By combining UHF with ultra-short TE, increased detection of small concentration metabolites is possible. This enables MRSI studies to acquire metabolite maps covering the full brain for low concentration metabolites which may be of particular interest in clinical studies.

In addition to the ultra-short TEs used in FID MRSI, it is common practice to utilize very short repetition times (TR) (Althaus et al., 2006; Dreher et al., 2003). By utilizing short TRs with elliptical shuttering, the acquisition time (TA) of MRSI data can be greatly reduced (Bogner et al., 2020). However, short TRs create strong T_1_-weighting of water and of metabolites. Previous studies (Bogner et al., 2012; Nassirpour et al., 2016) have reported good quality metabolite maps with short TR acquisitions; these studies report tissue contrast for total creatine (tCr), total choline (tCho), glutamate (Glu), glutamine (Gln), myo inositol (mI), and N-acetyl-aspartyl-glutamate (NAAG). However, measurements of T_1_-relaxation times would suggest that some of the observed contrast arises primarily from T_1_-weighting and not uniquely from concentration differences. For example, Glu T_1_-relaxation times estimated from human brain data at 9.4 T (Wright et al., 2021a) report a T_1_-relaxation time of approximately 1340 ms and 1470 ms for GM-rich and WM-rich regions respectively. Previous work (Gasparovic et al., 2006) proposed relaxation corrections for MRSI acquisitions using an internal water reference. In that work relaxation corrections were carried through for the water content of each voxel utilizing voxel-specific T_1_-relaxation times of water but only an averaged T_1_-relaxation time of metabolites.

In this study we expand this original T1 correction method (Gasparovic et al., 2006) by including voxel-tissue composition specific T_1_-corrections for all metabolites. Hence, we aim to correct T_1_-weighting in short TR ^1^H FID MRSI data and to report, for the first time, quantitative ^1^H MRSI results at 9.4 T in units of mmol kg^−1^. In the present work metabolite maps for 12 metabolites were acquired, and are showcased as T_1_-weighted and T_1_-corrected results. This study builds off of the previous work already performed at 9.4 T, by utilizing a high-resolution, single-slice FID MRSI acquisition (Nassirpour et al., 2016). While fitting metabolites, a simulated macromolecule (MM) spectrum (Wright et al., 2021a) was used to account for the MM contributes caused by using short TR, ^1^H FID MRSI. A simulated MM spectrum was utilized because experimentally acquiring the MM spectrum with a similarly short TR would is not possible to due to SAR intensive double inversion recovery pulses that would be used to null the metabolites (Nassirpour et al., 2017).

## 2. Methods

Experiments were performed on a 9.4 T whole-body scanner (Siemens Magnetom, Erlangen, Germany). A dual-row, 18Tx/32Rx phased array coil (Avdievich et al., 2018) was used to acquire data from ten healthy volunteers; however, due to poor data quality two volunteers were excluded from analysis; both subjects moved substantially during acquisitions of metabolite and water data. Thus, 8 healthy volunteers were included in the analysis (29 ± 2 years, 4 women, 4 men). A local ethics committee approved acquisition of data in human volunteers, and all volunteers provided written consent prior to scanning. The MRSI slice position for all volunteers was positioned with the bottom of the slice touching or slightly above the corpus callosum. A workflow diagram has been provided (Figure 1) to illustrate the processing and analysis pipeline as described in the following sections.

**Figure 1:**
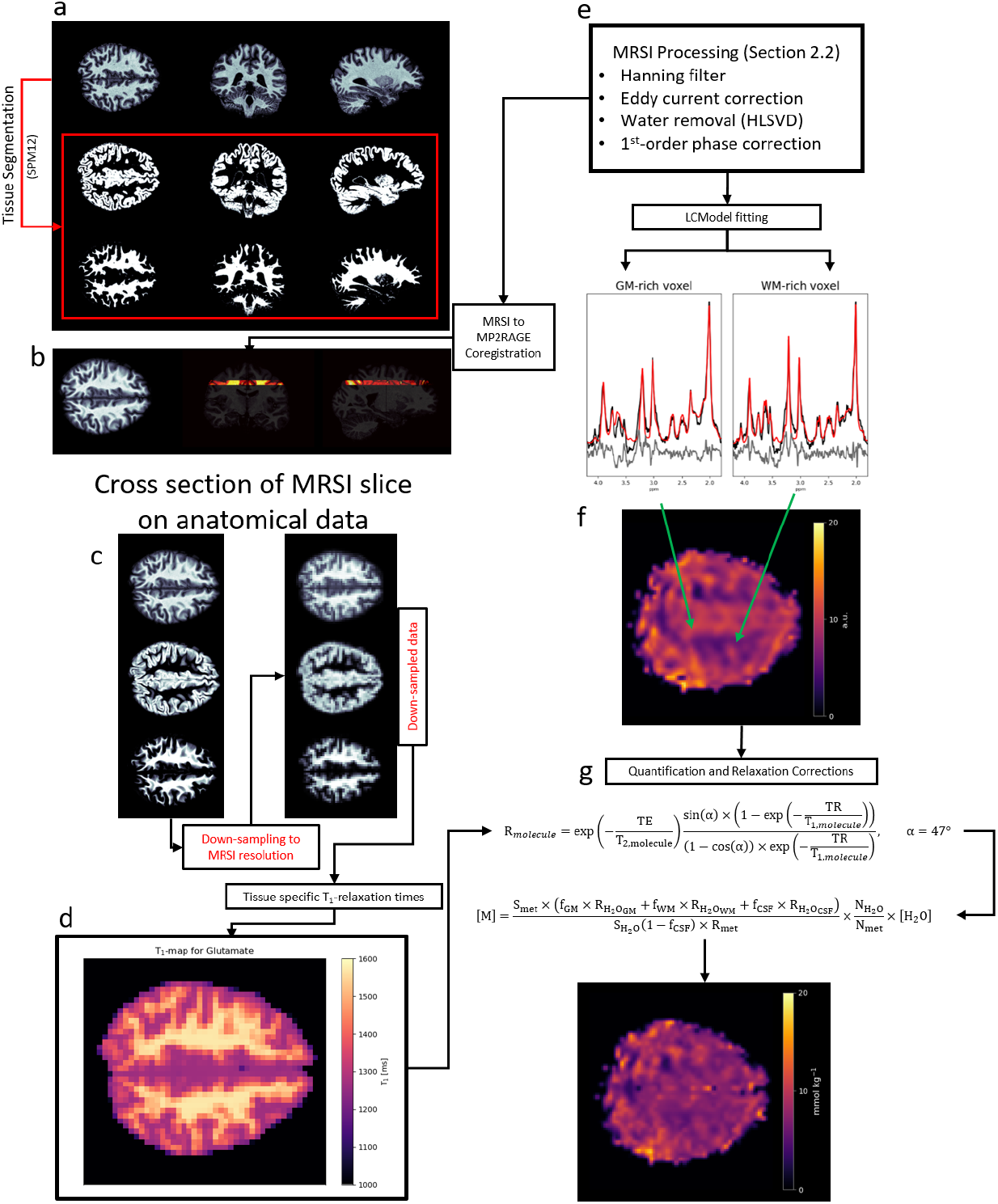
A workflow diagram as described in section 2 of the main text. (a) Anatomical data was segmented using SPM12, and then (b) coregistered to MRSI data. The coregistered anatomical data was then (c) down-sampled to match the MRSI resolution. (d) A T_1_-relaxation time map for each metabolite was then calculated and included in the quantification of each metabolite map in a voxel-specific manner. (e) _1_H MRSI data was preprocessed and (f) fitted using LCModel. (g) Water-normalized concentrations were corrected for partial volume effects, T_1_-weighting, and for contributing protons.

### 2.1 Data Acquisition

High-resolution (0.6 × 0.6 × 0.6 mm^3^) anatomical MP2RAGE images were acquired to be used for tissue segmentation. These images were acquired at the beginning of each scan session (TA = 11 min) with full details of the sequence described by (Hagberg et al., 2017) with a matched fast AFI map (Pohmann and Scheffler, 2013; Yarnykh, 2007) (resolution: 3.3 × 3.3 × 3.3 mm^3^, FA = 50°, TE/TR_1_/TR_2_ = 4/20/100 ms, TA = 1 min 20 sec) to correct for B_0_ distortions.

Spectroscopy data was acquired with a previously developed, elliptically shuttered, ^1^H FID MRSI sequence (Nassirpour et al., 2016) using a short TR (300 ms), acquisition delay, TE*, (1.5 ms), and flip angle (*α* = 47°). Water suppressed metabolite spectra utilized an optimized water suppression scheme with three unmodulated Hanning-filtered Gaussian pulses (BW = 180 Hz, duration = 5 ms) with flip angles of 90°, 79.5°, and 159°; the inter-pulse delay between all pulses was 20 ms. This study acquired ^1^H MRSI data with a 64×64 grid size at 210 × 210 × 8 mm^3^ (nominal voxel size = 3.28 × 3.28 × 8 mm^3^). In contrast to the previous work by (Nassirpour et al., 2016) where high-resolution metabolite mapping was performed and then receive sensitivity (B_1-_) corrected by a contrast-minimized GRE image, this study acquired a water reference ^1^H MRSI data set that matched water-suppressed acquisition parameters. The metabolite slice and water reference slice were acquired together in approximately 30 minutes. The water reference was used to account for B_1_ variations, eddy current corrections, and quantification. Vendor implemented, second-order shimming was performed for all data; the ^1^H MRSI metabolite data and water reference used the same shim settings.

### 2.2 MRSI Data Reconstruction and Processing

^1^H MRSI data were reconstructed using a spatial Hanning filter before the 2D FFT, then eddy current corrected which also corrects for zeroth-order phase (Klose, 1990), and coil combined using the singular value decomposition (SVD) method (Bydder et al., 2008). Following reconstruction, data post-processing included removing residual water using the Hankel-Lanczos (HLSVD) (Cabanes et al., 2001) method using 10 decaying sinusoids in the range of 4.4 to 5 ppm.

Data were then corrected for 1^st^-order phase distortion by means of linear back prediction of the missing FID points (Kay, 1988). Following back prediction, the phase of the metabolite data matched that of the basis set used in spectral fitting (Figure 1f). Data was then fitted as described in Section 2.4.

### 2.3 Anatomical Data Processing and MRSI Voxel Tissue Fractions

Anatomical data were reconstructed using an in-house developed MATLAB tool (Hagberg et al., 2017). Reconstructed data was then segmented into white-matter (WM), grey-matter (GM), and cerebrospinal fluid (CSF) probability maps using SPM12 (Figure 1a). The anatomical data and ^1^H MRSI data were aligned using an in-house developed python tool (Hunter, 2007; McKinney, 2010; Oliphant, 2006; van Rossum, 1995) to calculate the tissue fractions within each MRSI voxel. As shown in Figure 1a, the tissue probability maps and MRSI slice were coregistered (Figure 1b). The coregistered tissue probability maps were then resampled to the ^1^H MRSI resolution (3.28 × 3.28 × 8 mm^3^) (Figure 1c) (Van der Walt et al., 2014), and thus, all images showing anatomy in this work are a summation of the anatomical slices within the MRSI slice. The lower resolution tissue probability maps were then used to calculate T_1_-relaxation maps (Figure 1d) for T_1_-weighting corrections as described in Section 2.5.

### 2.4 Spectral Fitting

Spectra were fitted using LCModel (Provencher, 2001) from 1.8 to 4.2 ppm. Fit settings can be found in Supporting Information Annex A. A basis-set was simulated using the VeSPA suite (Soher, n.d.) for an FID sequence including 12 metabolites: NAA, tCr, aspartate (Asp), γ-aminobutyric acid (GABA), taurine (Tau), Gln, Glu, glutathione (GSH), mI, scyllo-inositol (Scyllo), NAAG, and phosphocholine + glycerophosphocholine (tCho). The MM spectrum was accounted for by using a sequence-specific simulated MM spectrum (MM_AXIOM_, (Wright et al., 2021a)). The MM_AXIOM_ used in this work assumed an equal contribution from WM and GM; (Wright et al., 2021b) showed that there was minimal benefit by introducing tissue specific MM spectra. Thus, to reduce the complexity in fitting, an average tissue composition was used for MM_AXIOM_ simulation. LCModel has the capability to fit MRSI data sets as a whole; however, data was fitted in LCModel as a batch of single voxel data as this yielded better spatial coverage. Following fitting in LCModel, data are referred to as water-normalized data. Data in this stage are still T_1_-weighted, but water-referenced.

### 2.5 T_1_-corrections

T_1_-corrected MRSI was originally shown by (Gasparovic et al., 2006); however, the relaxation corrections used in that study did not account for voxel-specific T_1_-relaxtion times of metabolites. In this work, we extend on the method described by (Gasparovic et al., 2006) by utilizing voxel-specific T_1_-corrections.

Previous studies at 9.4 T have reported the T_1_-relaxation times for water (Hagberg et al., 2017) and for metabolites (Deelchand et al., 2010; Wright et al., 2021a) in both GM and WM, and results suggest that T_1_-relaxation times vary by tissue compartment and not spatially. Thus, since the T_1_-relaxation phenomena is dependent primarily on tissue compartment and not on the spatial distribution or other factors, a linear relationship was used to estimate the T_1_-relaxation time of each metabolite for voxel-specific tissue fractions using T_1_-relaxation times from (Wright et al., 2021a).

Pure GM and WM T_1_-relaxation times of metabolites were used to calculate T_1_-relaxation maps as shown in Figure 1d. The T_1_-relaxation of each voxel was calculated by considering the T_1_-relaxation time to vary linearly depending on voxel tissue fraction:

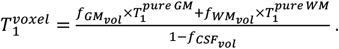

Where *f*(_*GM,WM*)*vol*_ is the GM and WM fraction within the voxel volume. Voxels with a *f*_*CSFvol*_ ≥ 30% were excluded from analysis.

### 2.6 Quantification

Following fitting of spectra in LCModel (Figure 1e), which resulted in water-normalized concentrations. These concentrations were corrected to be in mmolal (mmol kg^−1^) quantities. Partial volume effects as well as T_1_-weighting was corrected for during quantification. The relative densities of available MR-visible protons, a_y_, of water in GM, WM, and CSF were taken to be 0.78, 0.65, and 0.97, respectively

(Ernst et al., 1993). Relative densities were further scaled, 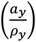, by the specific gravity of water for the tissue compartments (*ρ*_*GM*_, *ρ*_*WM*_, *ρ*_*CSF*_ = 1.04, 1.04, and 1.00 g/ml respectively (Brooks et al., 1980; Rieth et al., 1980; Torack et al., 1976)). The relaxation state of a metabolite, *R*_*met*_, and of water for a tissue compartment, *R*(_*H2O*)*y*_, are written as:

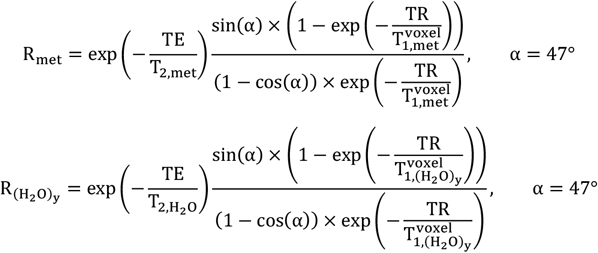

Where *α* is the flip angle applied, and 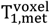 is the T_1_-relaxation time of a metabolite for a particular voxel. Calculation of the relaxation state (Figure 1g) excluded accounting for TE because of the short TE* used, and thus, the relatively small amount of T_2_-relaxation that could occur during that time. Following calculation of *R*_*met*_ and *R*(_H*2*O)y_, the concentration of a metabolite is calculated by:

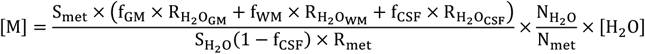

Where, 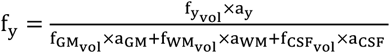, is the fraction of visible tissue volume; with y referring to the specific tissue compartment being calculated; note that a_y_ refers to the relative density scaled by the specific gravity, *ρ. S*_*met*_ and *S*_*H2O*_ are the signal of the metabolite and water respectively, and the concentration of water, [H_2_O], was assumed to be 55,510 mmol kg^−1^.

## 3. Results

### 3.1 MRSI Individual Spectra

LCModel fitting resulted in fitting of 12 metabolites (Figure 2). Residuals were minimal from 1.8-2.9 ppm, and broader residuals are present in varying amounts from 2.9-4.2 ppm; particularly for the left and right shoulders of the tCho resonance at 3.21 ppm. CRLB maps (Figure 4 and Figure 5) from LCModel show confident fitting for metabolites with sharp singlets (NAA, tCr, tCho, mI, and Glu) with the CRLB being less than 20 for NAA, tCr, Glu, mI, tCho, and Glx.

**Figure 2:**
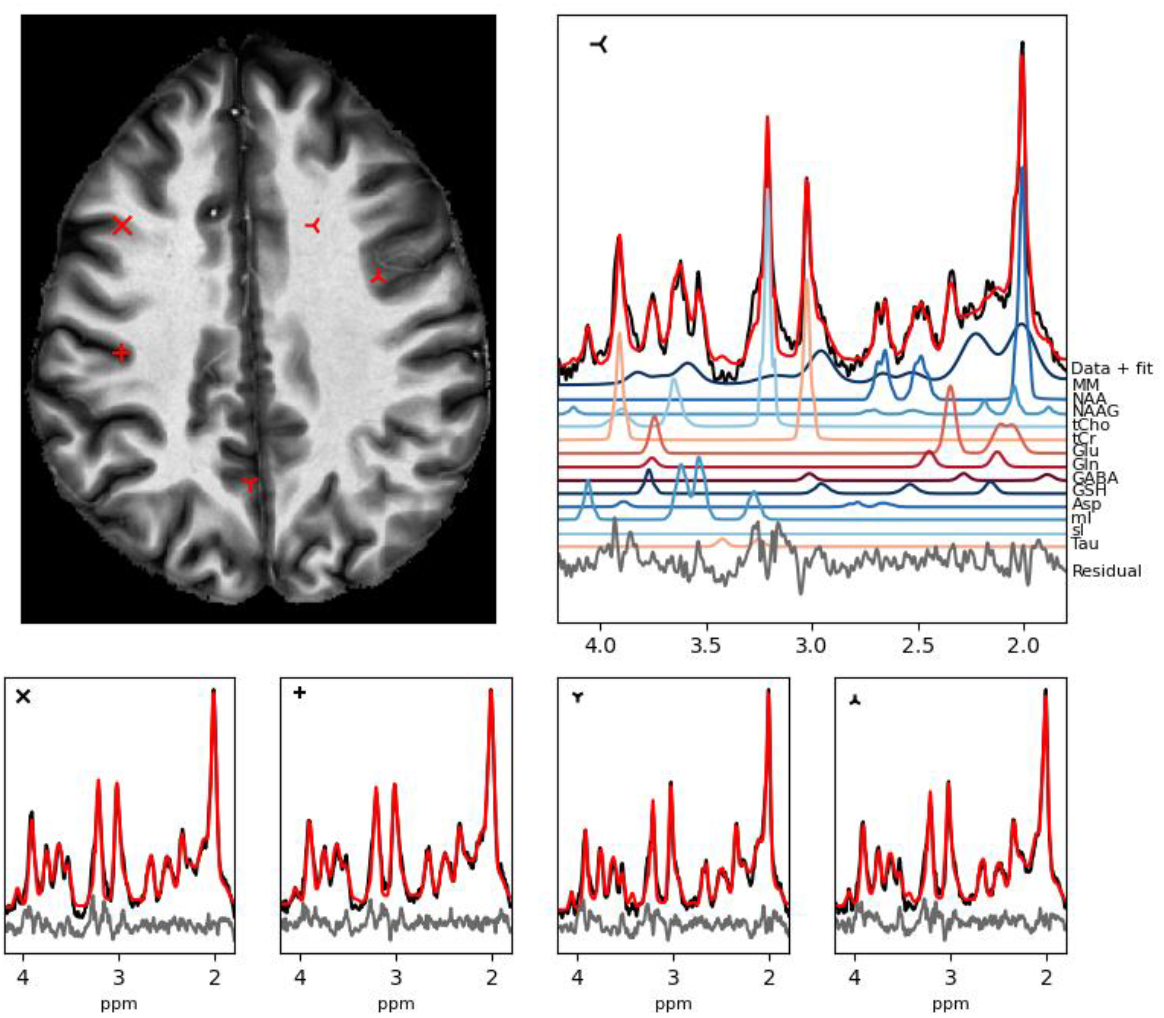
Representative spectra from voxels of various tissue composition. Metabolite spectra were fitted in LCModel using a basis set simulated using the VeSPA suite and an FID sequence. A simulated MM spectrum (MM_AXIOM_) was used to account for the MM contributions to the spectra. Markers in the top left of spectra correspond to matching locations on the MP2RAGE image.

**Figure 3:**
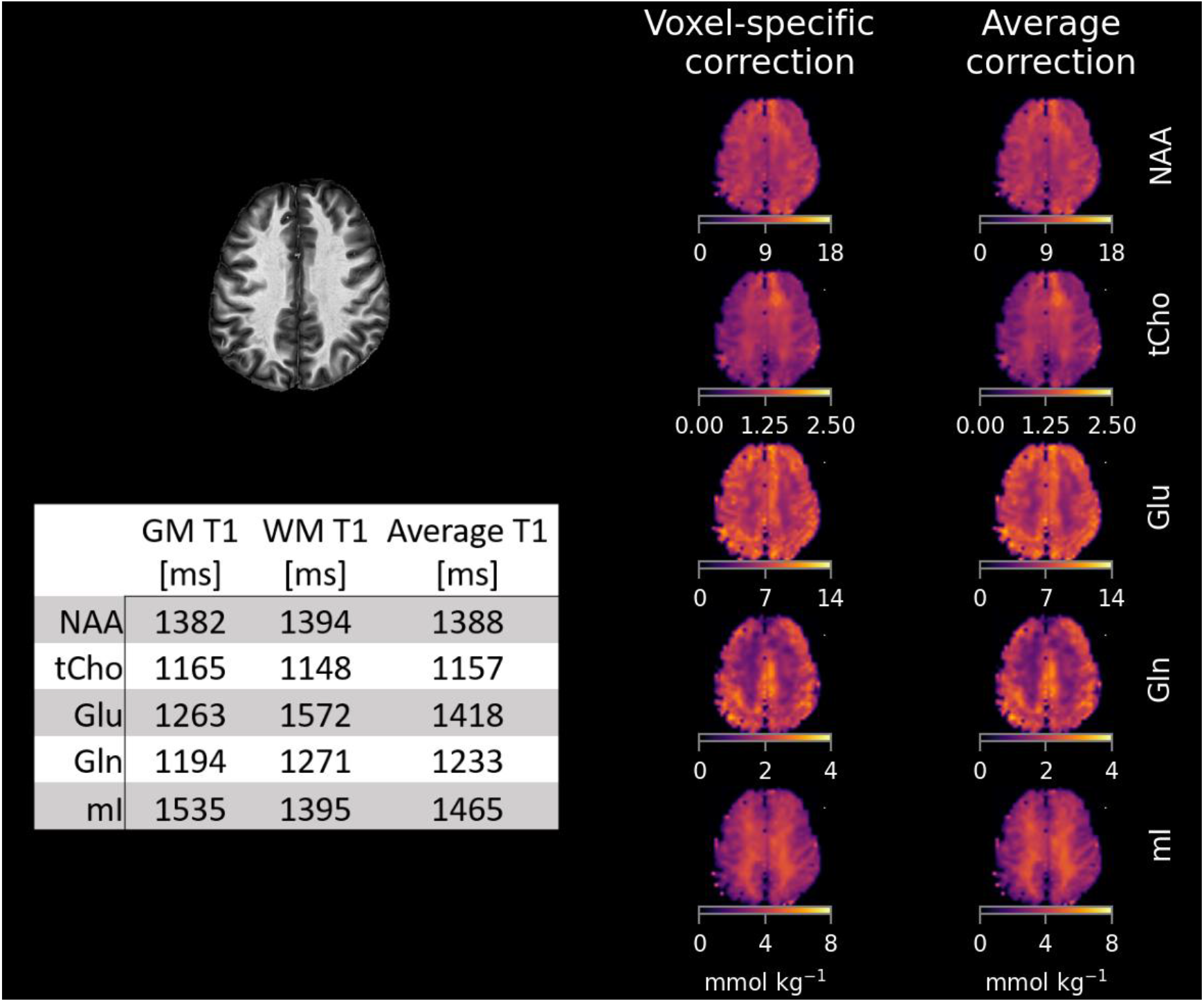
average T_1_-relaxation time correction (right column) and voxel-specific T_1_-corrections (left column). As can be seen, metabolite maps for metabolites that have similar T_1_-relaxation times in GM and WM (NAA, tCho) do not benefit as much with voxel-specific T_1_-corrections compared to metabolites with bigger differences (Glu, mI). A table of T_1_-relaxation times from 9.4 T (Wright et al., 2021a) is included as reference.

**Figure 4:**
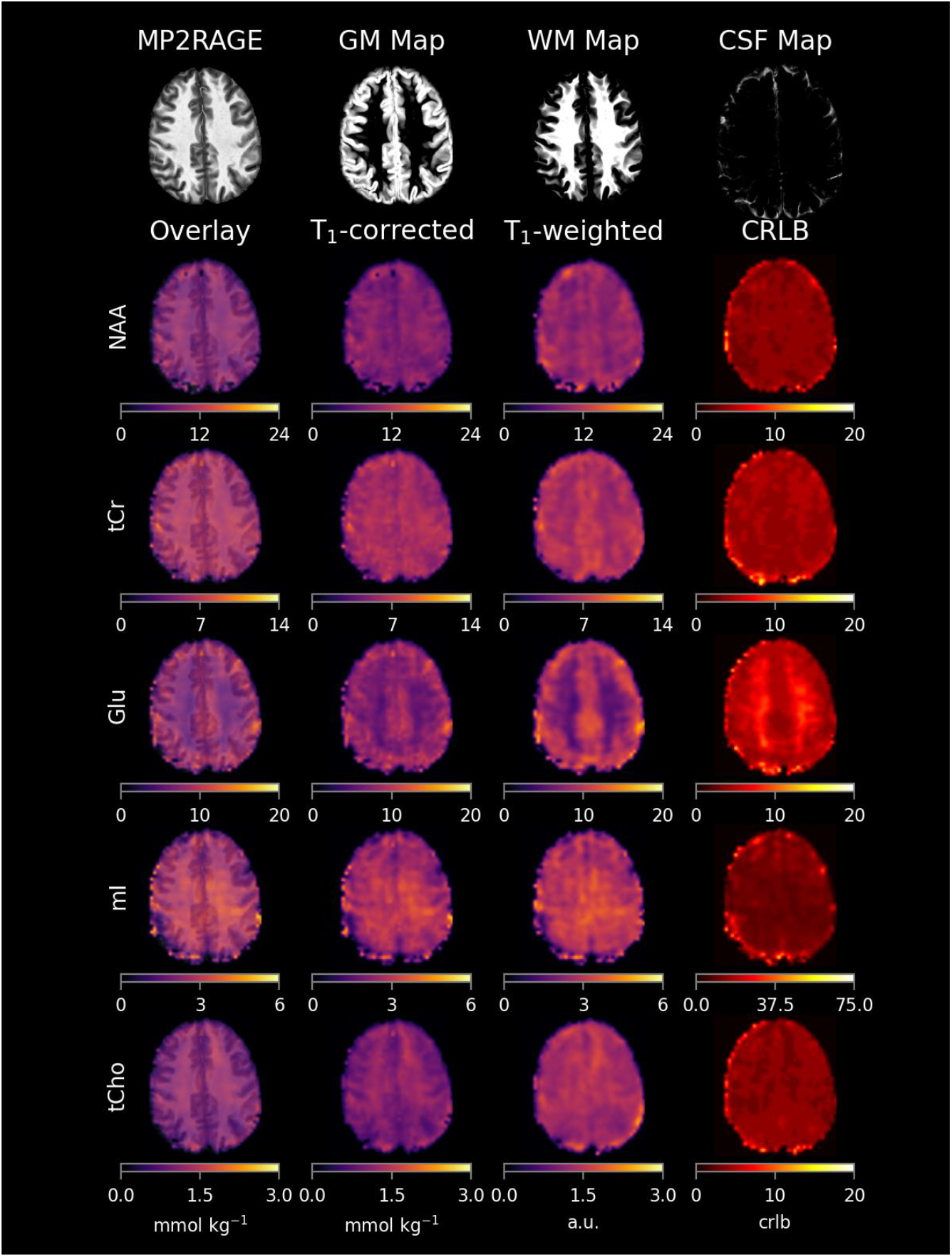
metabolite maps for NAA, tCr, Glu, mI, and tCho. Metabolite maps are showcased as being T_1_-weighted (as an LCModel output concentration), T_1_-corrected (quantitative units of mmol kg^−1^), and as Overlay (Quantitative maps overlaying MP2RAGE). Accompanying CRLB maps are in the far right column. Good anatomical detail is observed in all maps; with increased detail observed for T_1_-corrected metabolite maps. The total number of voxels included for all subjects are reported in Figure 7.

**Figure 5:**
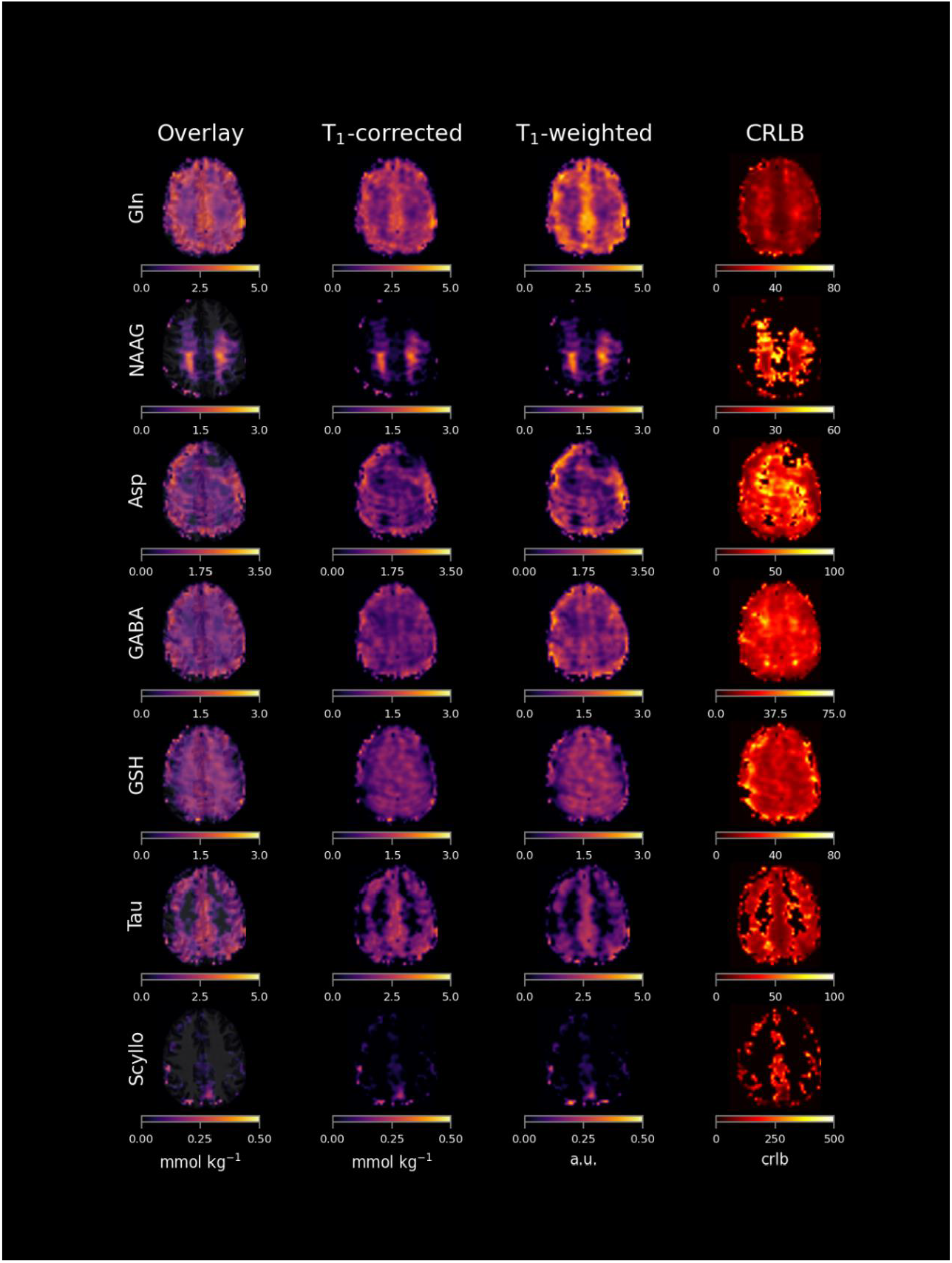
metabolite maps for Gln, NAAG, Asp, GABA, GSH, Tau. These metabolite maps have varying CRLB thresholds assigned based on data quality. Tissue dependence contrast is easily observed for Gln, NAAG, and Tau. GABA shows slight tissue contrast; however, T_1_-corrections reduce the contrast. The total number of voxels included for all subjects are reported in Figure 7.

The remaining metabolites (Gln, NAAG, Asp, GABA, GSH, Tau, and Scyllo) had different CRLB maxima chosen depending on the quality of the metabolite maps (Figure 5 and Figure 7). CRLB thresholds are given in Figure 4 and Figure 5 as the max value provided below the CRLB maps, and a table of CRLB thresholds is reported in Figure 7. The used CRLB thresholds were used for each metabolite individually, and thus, spectra were not entirely eliminated unless all metabolites were above the given CRLB threshold. CRLB thresholds were chosen to eliminate erroneous data (i.e. voxels with heavy lipid contamination where a metabolite was still fitted but not present) and not to set a rejection criterion for voxels as this could bias quantification results (Kreis, 2016).

### 3.2 Voxel-specific vs. average T_1_-corrections

A comparison of voxel-specific and average T_1_-corrections are shown in Figure 3. For metabolites with similar T_1_-relaxation times in GM and WM (i.e. NAA, tCho, and Gln), there is little observable difference between the methods. However, when the T_1_-relaxation times vary strongly between GM and WM there is a noticeable contrast shift in the metabolite maps. This observed change in contrast suggests that the strong GM/WM contrast for Glu can be attributed mainly to T_1_-relaxation time differences more than concentration differences. The differences observed in mI are marked most notably by a reduction in intensity in WM regions. Based on these findings, all metabolites in this work will utilize voxel-specific T_1_-relaxation times.

### 3.3 Metabolite Maps

Metabolite map comparisons for T_1_-weighted and T_1_-corrected data sets are showcased in Figure 4 and Figure 5 from the same subject. From left to right, columns in Figure 4 and 5 show the Overlay (T_1_-corrected map overlaying the MP2RAGE slices within the MRSI slice), T_1_-corrected maps (mmol kg^−1^), T_1_-weighted maps (a.u.), and CRLB maps. Metabolite maps show good spatial coverage of metabolite signal, encompassing most, if not all, of the anatomical slice. All maps in Figure 4 and Figure 5 included data, which had a CSF fraction less than 30 % as well as a CRLB less than the max CRLB value present in CRLB maps. The total number of voxels included for all subjects is presented in section 3.4.

T_1_-weighted metabolite maps agree well with recently reported metabolite maps from 9.4 T and 7 T (Nassirpour et al., 2016; Považan et al., 2017). By applying T_1_-correction in a voxel-specific manner (Figure 1g), we see that T_1_-corrections provide improved spatial specificity of ^1^H MRSI data. Furthermore, we observe diminished tissue contrast of certain metabolites (i.e. tCr and Glu) which previously had a stronger contrast with respect to signal intensity post-LCModel fitting (Figure 3 and Figure 4). Metabolites with similar T_1_-relaxation times in GM and WM exhibited less contrast change following T_1_-weighting corrections.

### 3.4 Metabolite Concentrations vs. Relative GM Fraction

To Illustrate the impact of partial volume effects and T_1_-weighting in short TR, ^1^H FID MRSI, Figure 6 shows linear regressions of Glu measurements against GM fraction and relative GM fraction, GM / (GM + WM), (A: Water-normalized Concentration without T_1_-correction vs GM fraction; B: Water-normalized Concentration without T_1_-correction vs Relative GM fraction; C: Concentration with T_1-_correction vs GM fraction; D: Concentration with T_1_-correction vs Relative GM fraction; A,B,C, and D: without CRLB or CSF exclusions). All data was pooled to visualize metabolite concentrations as a function of tissue fraction. The color bar present in all regression figures corresponds to the total tissue volume within a voxel (GM + WM). The GM fraction is a simple fraction showing the raw percentage of GM within a voxel, whereas the relative GM fraction is a metric to look at the amount of GM within a voxel without CSF contributions. In this work, we report the relative GM fraction as GM / (GM + WM); however, it can also be reported as GM / (1 – CSF) where CSF is the fractional CSF component.

**Figure 6:**
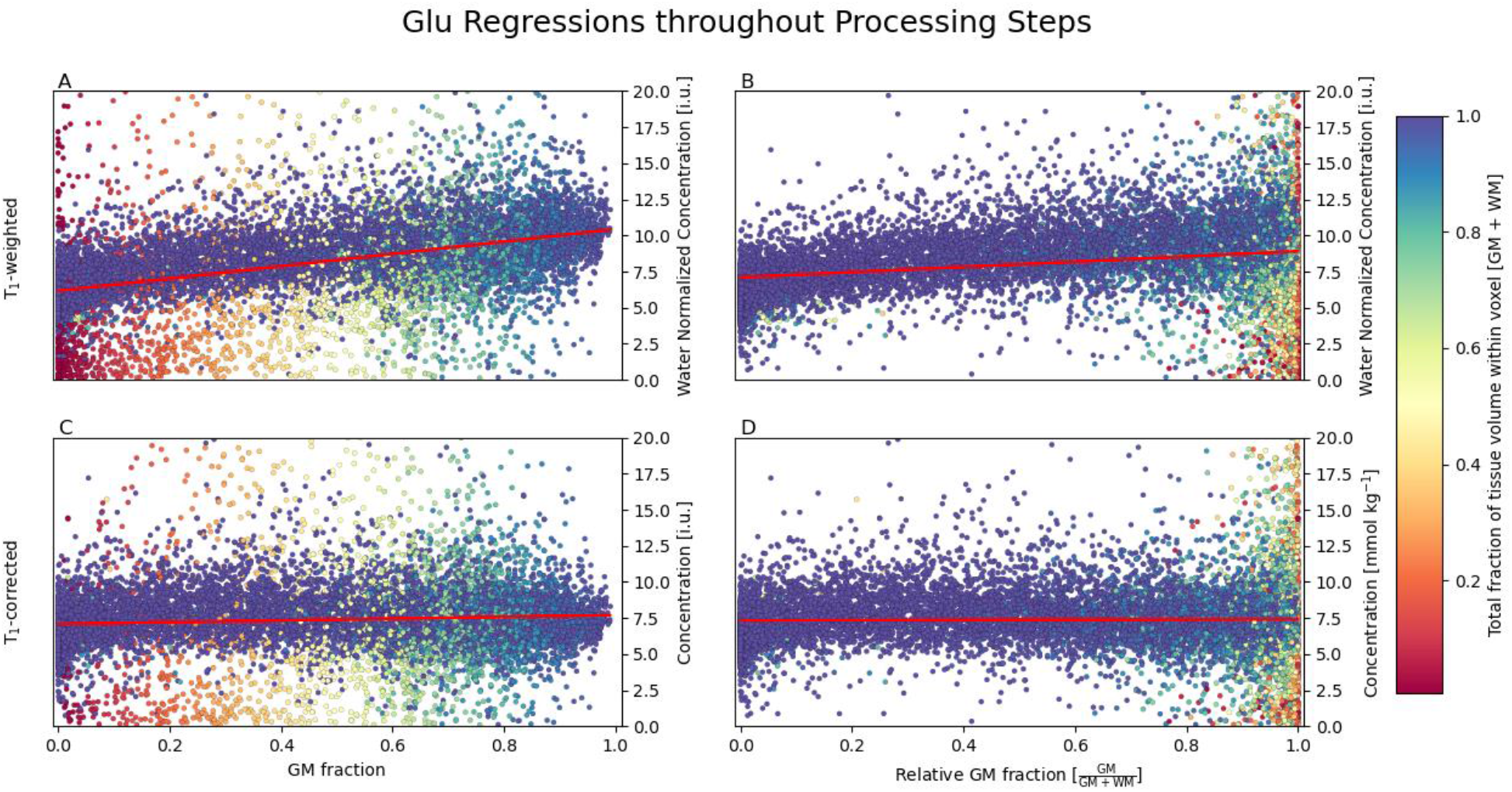
the impact of T_1_-corrections (top row to bottom row) and the impact of raw tissue fraction against relative tissue fraction (left column to right column) on data interpretation. The color bar shows the color of data points, and corresponds to the amount of tissue within a voxel (GM + WM). The red line is a linear fit to the data. All metabolite regressions are reported in Figure 7.

When comparing GM fraction to relative GM fraction, it is apparent that there are many voxels with a high relative fraction of GM but a low total tissue volume. The density of these data points shifts dramatically when comparing Figure 6 (A to B or C to D). The effect of the T_1_-correction is similar when comparing Figure 6 (A to C or B to D); for Glu, the tissue weighting that was apparent in Water-normalized concentrations is diminished. This is suggestive of there being no tissue contrast for Glu in the human brain.

Figure 7 shows T_1_-corrected concentrations of all metabolites regressed against relative GM fraction. Data from voxels that had less than 30% CSF content and with metabolite specific CRLB thresholds that were less than the CRLB value provided in the table in Figure 7 were included. The total number of voxels included into each regression is also reported in the table within Figure 7.

**Figure 7:**
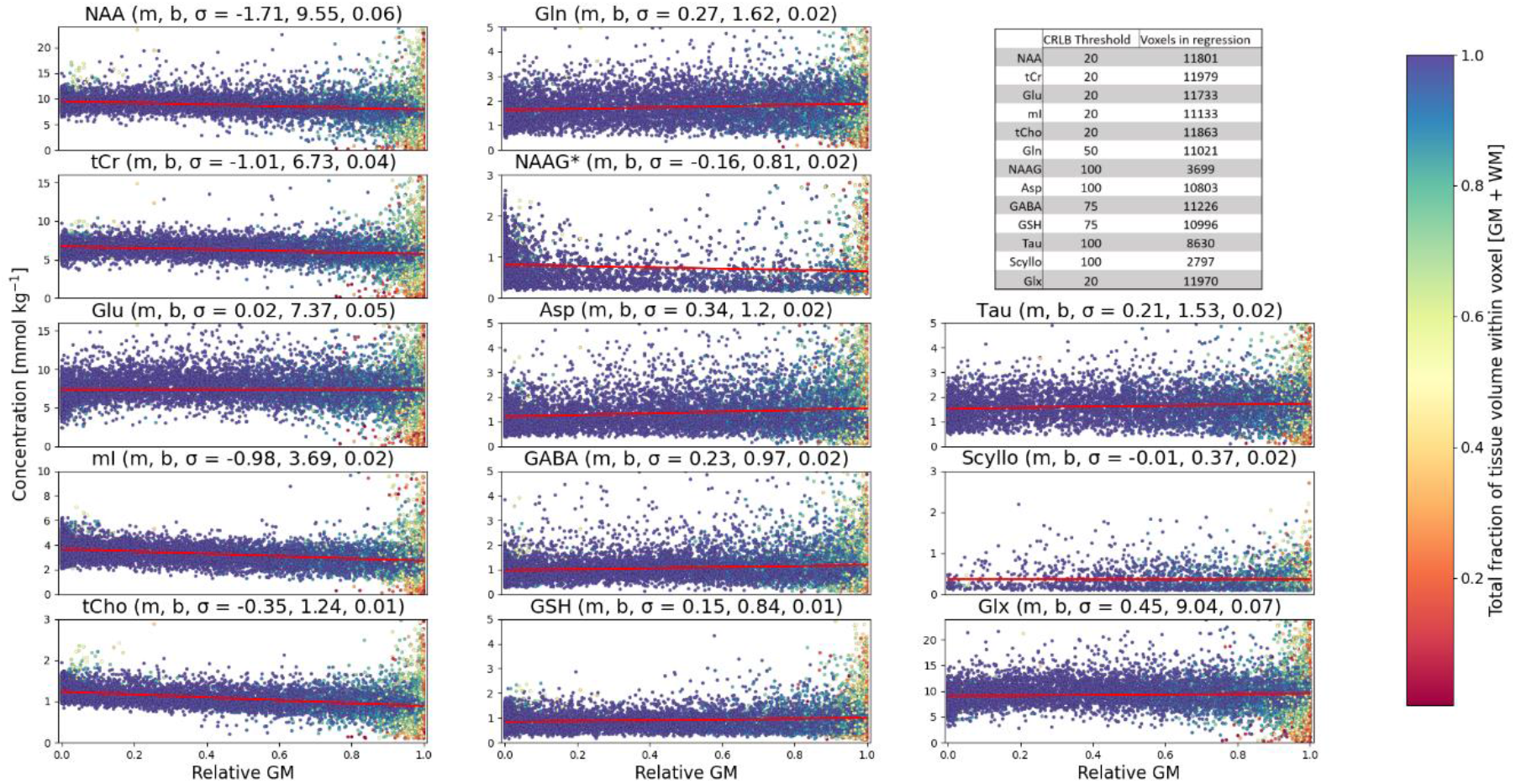
linear regressions for all metabolites are reported with respect to relative GM fractions [GM / (GM+WM)]. The color bar shows the color of data points, and corresponds to the amount of tissue within a voxel (GM + WM). Regressions were fitted with a standard straight line (y = mx + b); where m, b, and the standard error (σ) are reported next to the metabolite name. Tissue dependence can be estimated based on the slope of the regression fit line (red). The table in the top right shows the CRLB threshold for each metabolite (data with a CRLB over the given value were removed) and the total number of voxels included in the regression analysis. *Please refer to Supporting Information Figure S1 for NAAG regressions with respect to different relative GM fractions.

### 3.5 Metabolite quantification in mmol kg^−1^

Quantitative results (mmol kg^−1^) from this work are summarized and compared to previous studies in Table 1. To compare the results of this work to SVS studies, voxels were selected that were comprised of at least 70% GM or WM (GM-rich and WM-rich). Additionally, only voxels that had a minimum combined GM + WM fraction of 90% were included in concentration estimates. The total number of voxels counted for each metabolite are reported in Table 1. Discussion of this works results compared to other studies follow in section 4.1. Data from studies that reported results in mM (Kaiser et al., 2005; Pouwels et al., 1999; Pouwels and Frahm, 1998) are not included in Table 1, but are discussed as a point of reference in section 4.1. For further comparisons regarding the internal reference used, relaxation corrections, and corrections for macromolecules refer to Supporting Information Table 1.

**Table 1:** Quantitative results [mmol kg_-1_] from this work compared to results from SVS studies that acquired spectra in various brain regions. *GM-rich and WM-rich voxels were composed of a minimum of 70% of either GM or WM to be counted as a tissue-rich voxel; these voxels were also composed of a minimum total tissue fraction (GM + WM) of 90%. For further comparisons regarding the internal reference used, relaxation corrections, and corrections for macromolecules refer to Supporting Information Table 1.

| Quantitative results from 9.4 T <sup>1</sup>H MRSI and comparisons to SVS studies [mmol kg<sup>-1</sup>] |  |  |  |  |  |  |  |  |  |  |  |  |  |  |
| --- | --- | --- | --- | --- | --- | --- | --- | --- | --- | --- | --- | --- | --- | --- |
|  | Region | B<sub>0</sub><br>(T) | NAA | tCr | Glu | ml | tCho | Gln | NAAG | Asp | GABA | GSH | Tau | Scyllo |
| **This work** | GM-rich* | 9.4 | 8.1 ± 1.8 | 5.9 ± 1.1 | 7.6 ± 2.1 | 2.8 ± 1.0 | 0.9 ± 0.2 | 1.9 ± 0.7 | 0.9 ± 1.5 | 1.5 ± 0.9 | 1.1 ± 0.7 | 0.9 ± 0.4 | 1.7 ± 1.2 | 0.3 ± 0.3 |
|  | GM count |  | 2718 | 2728 | 2715 | 2559 | 2726 | 2633 | 308 | 2574 | 2614 | 2510 | 2296 | 917 |
| **This work** | WM-rich* | 9.4 | 9.3 ± 1.3 | 6.6 ± 0.8 | 7.2 ± 1.7 | 3.6 ± 0.7 | 1.2 ± 0.2 | 1.6 ± 0.6 | 0.9 ± 0.4 | 1.2 ± 0.6 | 1.0 ± 0.5 | 0.9 ± 0.3 | 1.6 ± 0.7 | 0.3 ± 0.3 |
|  | WM count |  | 3548 | 3561 | 3534 | 3522 | 3547 | 3453 | 1762 | 3086 | 3423 | 3473 | 2119 | 266 |
| **Deelchand et al., 2010** | occipital lobe | 9.4 | 13.5 ± 1.6 | 7.7 ± 0.4 | 9.3 ± 0.9 | 5.3 ± 0.4 | 0.9 ± 0.2 | 2.2 ± 0.2 | 1.1 ± 0.5 | 2.1 ± 0.5 | 1.3 ± 0.4 | 1.1 ± 0.3 | 1.3 ± 0.2 | 0.3 ± 0.2 |
| **Mangia et al., 2006** | occipital lobe | 7 | 10.8 ± 0.1 | 8.4 ± 0.3 | 11.0 ± 0.1 | 6.7 ± 0.1 | 1.3 ± 0.0 | 2.9 ± 0.1 | 1.4 ± 0.0 | 1.0 ± 0.2 | 1.0 ± 0.0 | 1.0 ± 0.1 | 1.9 ± 0.1 | 0.4 ± 0.0 |
| **Marjańska et al., 2012** | occipital lobe, mixed tissue | 7 | 12.6 ± 1.7 | 8.7 ± 1.1 | 9.6 ± 1.3 | 6.4 ± 0.8 | 0.9 ± 0.1 | 1.5 ± 0.5 | 0.4 ± 0.2 | 2.9 ± 0.8 | 1.5 ± 0.3 | 1.1 ± 0.1 |  | 0.4 ± 0.1 |
| **Marjańska et al., 2012** | motor cortex, WM-rich | 7 | 10.5 ± 1.2 | 7.4 ± 0.6 | 7.6 ± 0.9 | 5.9 ± 0.3 | 1.3 ± 0.1 | 1.0 ± 0.3 | 1.6 ± 0.4 | 2.6 ± 0.3 | 0.8 ± 0.6 | 0.7 ± 0.1 |  | 0.21 ± 0.1 |
| **Mekle et al., 2009** | occipital lobe, GM-rich | 7 | 11.8 ± 0.2 | 8.0 ± 0.4 | 9.9 ± 0.9 | 5.7 ± 0.5 | 1.1 ± 0.5 | 2.2 ± 0.4 | 1.1 ± 0.4 | 2.9 ± 0.5 | 1.3 ± 0.2 | 1.3 ± 0.2 | 1.5 ± 0.3 | 0.3 ± 0.1 |
| **Murali-Manohar et al., 2020** | occipital lobe, GM-rich | 9.4 | 12.6 ± 1.0 | 10.1 ± 0.5 | 10.9 ± 0.8 | 5.2 ± 0.5 | 1.0 ± 0.1 | 7.6 ± 0.9 | 1.4 ± 0.2 | 4.8 ± 1.2 | 1.9 ± 0.9 | 1.7 ± 0.2 | 1.6 ± 0.4 | 0.1 ± 0.1 |
| **Terpstra et al., 2009** | occipital lobe | 7 | 12.4 ± 0.7 | 8.4 ± 0.3 | 8.9 ± 0.3 |  | 1.0 ± 0.3 | 2.5 ± 0.2 | 1.2 ± 0.2 | 2.0 ± 0.4 | 0.9 ± 0.1 | 0.7 ± 0.1 | 1.7 ± 0.2 | 0.3 ± 0.1 |
| **Wright et al., 2021a** | occipital lobe, GM-rich | 9.4 | 12.5 ± 0.7 | 10.0 ± 1.1 | 13.0 ± 1.5 | 4.2 ± 0.8 | 1.2 ± 0.2 | 8.8 ± 1.1 | 1.8 ± 0.5 | 5.8 ± 0.6 | 1.9 ± 0.5 | 1.4 ± 0.3 | 1.5 ± 0.5 | 0.1 ± 0.05 |
| **Wright et al., 2021a** | occipital lobe, WM-rich | 9.4 | 11.8 ± 1.1 | 9.2 ± 0.8 | 10.9 ± 1.1 | 5.0 ± 0.6 | 1.8 ± 0.2 | 7.2 ± 1.1 | 2.6 ± 0.4 | 4.8 ± 0.7 | 1.2 ± 0.7 | 1.5 ± 0.3 | 1.0 ± 0.4 | 0.1 ± 0.1 |
| **Baker et al., 2008** | WM regions averaged | 3 | 17.6 ± 1.6 | 10.3 ± 1.2 |  | 5.1 ± 0.9 | 2.7 ± 0.5 |  |  |  |  |  |  |  |
| **Baker et al., 2008** | GM regions averaged | 3 | 15.7 ± 1.5 | 11.3 ± 1.4 |  | 5.7 ± 1.0 | 1.8 ± 0.4 |  |  |  |  |  |  |  |
| **Michaelis et al., 1993** | parietal lobe, GM-rich | 2 | 9.8 ± 1.4 | 6.6 ± 1.0 | 9.3 ± 2.1 | 4.7 ± 0.8 | 1.1 ± 0.3 |  | 0.0 ± 0.0 |  |  |  |  | 0.4 ± 0.1 |
| **Michaelis et al., 1993** | parietal lobe, WM-rich | 2 | 7.8 ± 0.7 | 5.3 ± 0.7 | 6.6 ± 1.2 | 3.9 ± 0.9 | 1.6 ± 0.3 |  | 2.2 ± 0.8 |  |  |  |  | 0.3 ± 0.1 |

## 4. Discussion

### 4.1 Metabolite maps and quantification of ^1^H MRSI data

The following sections discuss metabolite maps and quantification results from the 12 metabolites reported in this study. The spectral quality and the spatial coverage of acquired ^1^H FID MRSI data allows us to observe metabolite signals across the full anatomical slice for 12 metabolites. Holes throughout metabolite maps arise either from a high CSF fraction (≥ 30%) or from a CRLB greater than the max value reported in the CRLB table of Figure 7. The total number of fitted voxels for the whole study is also reported in Figure 7.

In line with previous work, this study confirms that metabolite mapping improves dramatically at UHF. While metabolites such as NAA, tCr, tCho, Glx, and mI can be imaged at field strengths below 7 T, results from this work and previous ^1^H MRSI data obtained at UHF (Bogner et al., 2012; Hangel et al., 2021;

Henning et al., 2009; Nassirpour et al., 2016; Považan et al., 2017) show substantial improvements with respect to the spatial coverage and spatial resolution of respective metabolite maps. The metabolite maps shown in Figure 4 are in general agreement with those from previous UHF ^1^H MRSI studies (Bogner et al., 2012; Hangel et al., 2021; Henning et al., 2009; Nassirpour et al., 2016).

The further benefit that comes from UHF is being able to reliably image metabolites with lower concentrations or those that are overlapped by other metabolite peaks with stronger signal intensity at lower field strengths (i.e. Glu, Gln, NAAG, and Tau). Figure 5 shows metabolite maps for metabolites with lower concentrations or reduced fitting accuracy. While these metabolites can be challenging to fit with ^1^H MRSI acquisitions we still see good spatial resolution of many of these metabolites. UHF strengths lend themselves to detecting more metabolites, and within the realm of UHF, 9.4 T has improved detection capabilities compared to 7 T.

In general, single slice, quantitative metabolite maps (Figure 4 and Figure 5) as well as pooled data from regressions (Figure 7) show metabolite concentrations comparable to reported SVS studies (Baker et al., 2008; Deelchand et al., 2010; Kaiser et al., 2005; Mangia et al., 2006; Marjańska et al., 2012; Mekle et al., 2009; Michaelis et al., 1993; Murali-Manohar et al., 2020; Pouwels et al., 1999; Pouwels and Frahm, 1998; Terpstra et al., 2009; Wright et al., 2021a). To maintain consistency, all comparisons in Table 1 were performed with studies that also reported concentrations in units of mmol kg^−1^. Linear regressions (Figure 7) show metabolite concentrations as a function of relative GM fraction. For detailed comparisons of concentrations, please refer to Table 1. In the following sections, anatomical detail and tissue contrast are used to describe the quality of metabolite maps. Anatomical detail refers to seeing fine details such as GM folding, whereas tissue contrast refers to the in general contrast of intensity between GM and WM regions.

#### 4.1.1 NAA

NAA maps show slight anatomical detail in T_1_-weighted metabolite maps (Figure 4), and this anatomical detail is improved in T_1_-corrected maps. NAA has a concentration that is slightly lower than those in previous GM-rich reports (Deelchand et al., 2010; Mangia et al., 2006; Marjańska et al., 2012; Mekle et al., 2009; Murali-Manohar et al., 2020; Terpstra et al., 2009; Wright et al., 2021a); however, when considering the NAA/tCr ratio the quantification results are similar. NAA concentrations are similar for previous WM-rich reports (Baker et al., 2008; Pouwels et al., 1999; Pouwels and Frahm, 1998), although these studies report results in mM quantities. A further improvement to NAA quantification is discussed in section 4.3.

#### 4.1.2 tCr

T_1_-weighted tCr maps show a slight GM/WM tissue contrast; however, this contrast is less observable following T_1_-corrections. While the GM/WM contrast is diminished with T_1_-corrections, the anatomical detail is improved.

tCr concentrations range from approximately 6-7 mmol kg^−1^ for GM-rich and WM-rich voxels respectively. tCr concentrations from this work are in general lower than previous reports; many factors can influence these differences (i.e. relaxation corrections, concentration assumptions, and basis sets used in fitting in combination with acquisition sequences). Therefore, when discussing similarities of other metabolite concentrations, it is important to consider systematic differences of the tCr concentration. The need for this comes from how LCModel fitting utilizes the tCr peak at 3.03 ppm to calculate water-scaled concentrations, and thus, by assigning a concentration to tCr, the other metabolites will be scaled by this concentration. A further improvement to tCr quantification is discussed in section 4.3.

#### 4.1.3 Glu

Glu maps display good anatomical detail in both T_1_-weighted maps and T_1_-corrected maps; however, the contrast in Glu concentration between GM and WM is reduced in T_1_-corrected maps. This is due to the longer T_1_-relaxation time of Glu in WM (Wright et al., 2021a; Xin et al., 2013), and thus, the relaxation correction term, R_Glu_, is less in WM and results in an increase in estimation of Glu concentrations in WM. While metabolite maps still show contrast between GM and WM for T_1_-corrected Glu concentrations; regressions (Figure 7) suggest that there is little difference between Glu concentrations in GM and WM. The Glu concentrations are in a similar range to previous studies listed in Table 1; however, in general are lower in concentration.

T_1_-weighted metabolite maps in this work show strong tissue contrast, which is consistent with previous literature showing increased concentration of Glu in GM (Bogner et al., 2012; Henning et al., 2009; Nassirpour et al., 2016; Považan et al., 2017). However, this contrast is greatly diminished when performing voxel-specific T_1_-relaxation corrections (Figure 3 and Figure 4).

#### 4.1.4 mI

mI quantification reveals decreased concentrations compared to other works when considering GM-rich voxels (approx. 2.8 mmol kg^−1^, Table 1). mI maps show increased concentration of mI in WM-voxels. The increased WM contrast is also better observed following voxel-specific T_1_-corrections which result in improved anatomical detail across the MRSI slice. The concentration of mI is approximately 3.6 mmol kg^−1^ in WM-rich voxels. This concentration is lower than previous reports; however, previous work which investigated the use of a MM_AXIOM_ (Wright et al., 2021a, 2021b) has shown that overestimations of mI are common when using T_1_-weighted MM spectra for fitting.

Previous works at UHF that did not incorporate simulated MM spectra to account for contaminations from MM showed the mI distribution to be more localized in GM (Bogner et al., 2012; Henning et al., 2009; Nassirpour et al., 2016). mI maps presented in Figure 3 and Figure 4 display higher concentrations of mI in WM than in GM. Increased mI concentrations can be traced acrossed WM with decreasing mI concentrations apparent in regions at the center and periphery of the slice with increased GM tissue fractions and GM folding. This finding is in agreement with 7 T work which also used simulated MM spectra in fitting (Považan et al., 2017) that showed a higher concentration of mI in the WM. Furthermore, this is supported by CEST findings (M et al., 2011) of mI having higher concentrations in human brain WM.

#### 4.1.5 tCho

In T_1_-corrected maps, tCho shows higher concentrations in WM, and tCho maps show excellent anatomical detail. In addition to the apparent GM/WM contrast, tCho also looks to have higher concentrations toward the frontal lobe, which is consistent with previous literature (Bogner et al., 2012; Henning et al., 2009; Henning, 2017; Maudsley et al., 2009; Nassirpour et al., 2016; Považan et al., 2017).

tCho concentrations from this work agree well with SVS studies for GM-rich and WM-rich investigations (Baker et al., 2008; Deelchand et al., 2010; Kaiser et al., 2005; Mangia et al., 2006; Marjańska et al., 2012; Mekle et al., 2009; Michaelis et al., 1993; Murali-Manohar et al., 2020; Pouwels et al., 1999; Pouwels and Frahm, 1998; Terpstra et al., 2009; Wright et al., 2021a). tCho concentrations range from 1.2 – 0.9 mmol kg^−1^ when going from WM-rich to GM-rich voxels. Another consideration for tCho would be to do regional comparisons or spatial distribution since tCho has been reportedly higher in concentration in the frontal cortex compared to posterior brain regions (Hangel et al., 2021). The recent work by (Hangel et al., 2021) highlights this well by having multiple slices spanning the cerebrum.

The tCho resonance at 3.2 ppm displays increased residuals at the shoulders of potentially due to the broadness of the lineshapes of FID-MRSI data. In fitting trials, there was an attempt to include accompanying resonances (i.e. phosphoryl ethanolamine and phosphocholine); however, consistent and reliable fitting was not achieved when using combinations of the aforementioned two metabolites in addition to the GPC resonances.

#### 4.1.6 Gln

Tissue contrast is still observed for Gln following voxel-specific T_1_-corrections. This may suggest that the concentration differences reported for Glx (Baker et al., 2008; Hangel et al., 2021, 2018) between GM and WM can be attributed primarily to Gln. Metabolite maps (Figure 5) for Gln are consistent with previous reports at 9.4 T (Nassirpour et al., 2016). While the contrast intensity is slightly reduced in comparison to the previously referred results for T_1_-corrected metabolite maps, the general GM/WM contrast is still well observed. Indeed Gln maps from this work are in agreement with previous 9.4 T results (Nassirpour et al., 2016), and they also show improvement in detection. The present results show much improved fitting of Gln across the full slice. The improved detection capability of Gln, along with quantitative results at UHF, provides a means to investigate Gln more precisely. Gln concentrations are in line with most previous literature (Table 1); excluding 9.4 T results which utilized semiLASER acquisitions (Table 1).

#### 4.1.7 NAAG

NAAG maps show that NAAG is almost exclusively located in WM. This is consistent with multiple studies from 9.4 T (Nassirpour et al., 2016; Wright et al., 2021a) and 7 T (Henning et al., 2009; Považan et al., 2017; Xin et al., 2013). The studies estimating T_1_-relaxation times (Wright et al., 2021a; Xin et al., 2013) both show a large increase in standard deviation for NAAG T_1_-relaxation time estimates for GM-rich voxels compared to their WM-rich counterparts. NAAG concentrations are in a similar range to most previous literature (Table 1); excluding 9.4 T results which utilized semiLASER acquisitions (Table 1).

When considering the regressions of NAAG (Figure 7), it is reasonable to come to the conclusion that a regression from a relative GM fraction of 0 to 1 for NAAG concentrations is unreasonable. Supporting Information Figure S1 shows NAAG concentrations regressed with varying relative GM fraction thresholds CRLB thresholds. These regressions support the observations from metabolite maps (Figure 5) that NAAG is located primarily in WM.

#### 4.1.8 Asp

Asp maps highlight the difficulty in fitting Asp. However, it can be ascertained from these maps that Asp has a more prominent distribution in GM-rich regions due to the increased CRLBs in WM-rich regions.

The regression of Asp in Figure 7 shows a slight positive slope; indicating an increased Asp concentration in GM-rich regions. Previous work at 9.4 T (Nassirpour et al., 2016) report a similar distribution of Asp in the human brain, with a similar slice position. Asp concentrations for GM-and WM-rich regions (Table 1) are similar to previous reports from 7 T, and lower in concentration in comparison to previous 9.4 T results (with the exception of the work from (Deelchand et al., 2010)). The works at 9.4 T which utilized semiLASER (Murali-Manohar et al., 2020; Wright et al., 2021a) highlighted the observation of increased Asp results to other literature.

#### 4.1.9 GABA

GABA remains a challenging metabolite to fit in the standard (non-edited) metabolite ^1^H spectrum at 9.4 T. GABA metabolite maps show some anatomical detail, and from T_1_-weighted maps (Figure 5) there appears to be an increased concentration in GM. This contrast is consistently observed in T_1_-weighted data that spans a full slice (Moser et al., 2019; Nassirpour et al., 2016). However, T_1_-corrected metabolite maps reduce this contrast. Meanwhile, the GABA regression in Figure 7 shows a slight elevation in GM compared to WM. Future work would benefit by improving the fitting of GABA in non-edited metabolite spectra.

GABA concentrations in GM-rich regions agree with those published in previous studies with being approximately 1 mmol kg^−1^ (Deelchand et al., 2010; Mangia et al., 2006; Marjańska et al., 2012; Mekle et al., 2009; Murali-Manohar et al., 2020; Terpstra et al., 2009; Wright et al., 2021a). GABA remains a difficult metabolite to fit at 9.4 T, and thus, more advanced methods such as spectral editing techniques are likely better served toward quantifying GABA. From a large cohort, cross-site, and cross-vendor study of GABA concentrations using a MEGA-PRESS technique, the estimated concentration of GABA+ (GABA + MM contamination) was found to be approximately 0.116 (GABA/tCr). This result is similar within reason to GM- and WM-rich regions in this work, approximately 0.18 and 0.15, respectively.

#### 4.1.10 GSH

GSH was detected in most voxels, and quantified results are in agreement with previous GM-rich studies (Deelchand et al., 2010; Mangia et al., 2006; Marjańska et al., 2012; Murali-Manohar et al., 2020; Terpstra et al., 2009; Wright et al., 2021a). GSH metabolite maps show that GSH was fit across the full slice, but maps do not show a clear contrast between GM and WM (Figure 5). While the regression in Figure 7 is suggestive of a slight increase in concentration for GM-rich regions. Concentration estimates from GM- and WM-rich results show very similar results for both tissue types. However, at the periphery of the brain, GSH was difficult to fit potentially due to increased linewidths, and this could cause the discrepancy observed compare to previously 9.4 T GSH maps (Nassirpour et al., 2016).

#### 4.1.11 Tau

Tau shows opposite contrast to NAAG, and is located almost exclusively in GM-rich voxels. Tau metabolite maps match the tissue contrast reported from previous ultra-high field MRSI studies (Bogner et al., 2012; Nassirpour et al., 2016; Považan et al., 2017). The concentration of Tau in GM-rich voxels is approximately 1.7 mmol kg^−1^. This is consistent with reports from GM-rich studies (Table 1). While the WM voxel count is similar to the GM voxel count, the distinct GM localization can be realized when looking at the difference in GM and WM voxel counts. It is apparent that Tau is still fitted in WM-rich voxels, and this could be due to the thru-plane CSDE that occurs from the 2D slice excitation.

#### 4.1.12 Scyllo

Scyllo was not well fit in this study. Only 2797 voxels were fitted out of all of the voxels; this is very few compared to most of the other metabolites having approximately 10000 useable voxels. Given that Scyllo is also difficult to fit in SVS studies at 9.4 T (Murali-Manohar et al., 2020; Wright et al., 2021a), it is reasonable that Scyllo is difficult to fit with short TR, ^1^H FID MRSI acquisitions. However, the limited fitting of Scyllo is consistent with previous 9.4 T results (Nassirpour et al., 2016); thus, it could be that Scyllo is located in the GM in very low concentrations. Future work could investigate Scyllo distributions using MRSI techniques with more averages, or by utilizing SVS localization with voxels almost exclusively located in GM and WM.

### 4.2 Voxel-specific vs Average T_1_-corrections

Foundational work from (Gasparovic et al., 2006) reported using an internal water reference and relaxation corrections for ^1^H MRSI data using an average T_1_-relaxation time for each metabolite, and many studies have followed this practice since. However, in this work we utilized voxel-specific T_1_-corrections for each metabolite. Both methods utilize the tissue specific relaxation time of water measured at 9.4 T (T_1_-relaxation time in GM/WM/CSF = 2120/1400/4800 ms (Hagberg et al., 2017)). As shown in Figure 3, the difference between average T_1_-corrections and voxel-specific T_1_-corrections is insubstantial for many metabolites. When the difference in T_1_-relaxation time is minimal, the metabolite maps appear nearly identical for both voxel-specific and average T_1_ correction methods.

However, for metabolites such as Glu or mI the difference in T_1_-relaxation leads to slight differences in metabolite maps between the two methods. Glu maps with voxel-specific T_1_-corrections show an increased intensity in WM voxels; which manifests as a slight reduction in GM/WM contrast. Alternatively to Glu, Gln has a similar T_1_-relaxation time in GM and WM (approx. 1200 ms (Wright et al., 2021a)), and for Gln maps (Figure 5), it is apparent that tissue contrast is slightly reduced following T_1_-corrections.

mI maps (Figure 3 and Figure 4) also benefit from voxel-specific T_1_-corrections.The contrast between GM and WM is improved following voxel-specific T_1_-corrections as well as some increased evidence of anatomical detail seen as better separation between GM and WM both in the center and periphery of the MRSI slice.

The T_1_-relaxation times used from 9.4 T measurements (Wright et al., 2021a) are extrapolated to pure-GM and pure-WM times. The extrapolated times show increased standard error, and thus, by can contribute to slight errors in metabolite quantification. Future work investigating T_1_-relaxation times across a more diverse sample of tissue densities would benefit voxel-specific T_1_-corrections. One approach to estimate these times could be to perform a saturation recovery method for an MRSI slice, and apply the measured T_1_-relaxation times directly to acquired voxels.

### 4.3 Advancing quantitative MRSI in the future

Voxel-specific T_1_-corrections improved the overall accuracy of metabolite maps. Nonetheless, there is room for further improvement of T_1_-corrections by splitting the CH_2_ and CH_3_ moieties (i.e. for NAA and tCr). These moieties have different T_1_-relaxation times as reported by (Deelchand et al., 2010; Wright et al., 2021a; Xin et al., 2013). In addition to the difference in T_1_-relaxation times for metabolite moieties, some moieties have large differences between GM and WM (i.e. NAA-CH3 and tCr-CH3). This could contribute to the tissue contrast observed for tCr, and future work would benefit from exploring metabolite mapping when fitting short TR, ^1^H FID MRSI with split metabolite moieties while using an MM_AXIOM_. Furthermore, applying moiety specific T_1_-relaxation corrections for NAA and tCr could likely improve the concentration estimate and align the results more directly to the SVS results used as a comparison in Table 1.

While there are a variety of methods to further accelerate MRSI acquisitions (Bogner et al., 2020) this work acquired fully sampled ^1^H MRSI data. This work aims to provide a reference for quantitative data at UHF; with this study, other studies should be able to directly compare quantitative results acquired with accelerated techniques. This reference can provide valuable insight into how to utilize acceleration techniques such as highly-accelerated water referencing (Chang et al., 2018) or variable density under-sampling techniques (Nassirpour et al., 2018b, 2018a) for metabolite quantification, and therefore, make quantitative metabolic imaging with ^1^H MRSI more clinically feasible.

## 5. Conclusion

This work reports the first quantitative ^1^H MRSI results acquired at 9.4 T in the human brain. To quantify fully sampled ^1^H FID MRSI, an internal water reference was used and voxel-specific T_1_-corrections were performed to account for the T_1_-relaxation time differences apparent for metabolites between GM- and WM-rich regions. Voxel-specific T_1_-corrections show that when not accounting for a strong difference in T_1_-relaxation times results may be biased and show increased contrast. By utilizing a MM_AXIOM_ and voxel-specific T_1_-relaxation times, this work was able to present high resolution, quantitative metabolite maps for 12 metabolites. Quantified metabolites included NAAG, Gln, GABA and Tau, which have been notoriously challenging metabolites to fit. Applying voxel-specific T_1_-corrections to heavily T_1_-weighted MRSI data allows for direct comparison to many other studies, and thus, brings ^1^H FID MRSI a step closer to expanded clinical application.

## Acknowledgements

The authors are appreciative of helpful conversations from Tamas Borbath, Johanna Dorst, Kye Stachowski, Loreen Ruhm, and Theresia Ziegs.

## Notes

### Competing Interest Statement

The authors have declared no competing interest.

