## Supporting Information for "Quantitative T_1_-relaxation corrected metabolite mapping of 12 metabolites in the human brain at 9.4 T"

Supporting Information Figure 1: NAAG fitting at relative GM thresholds of 60, 40, and 20 % (top, middle bottom). Various CRLB thresholds are also included to show NAAG how NAAG fit confidence impacts regression performance. CRLB = 100 is equal to that listed in Figure 7.

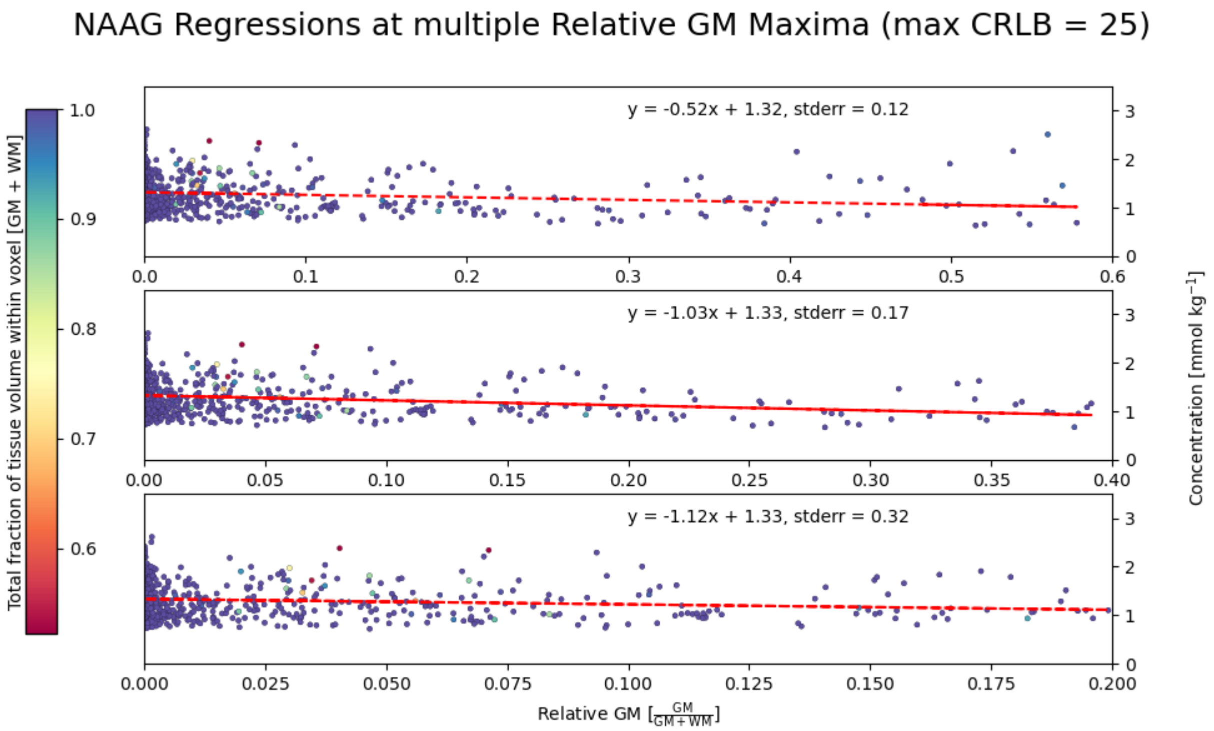


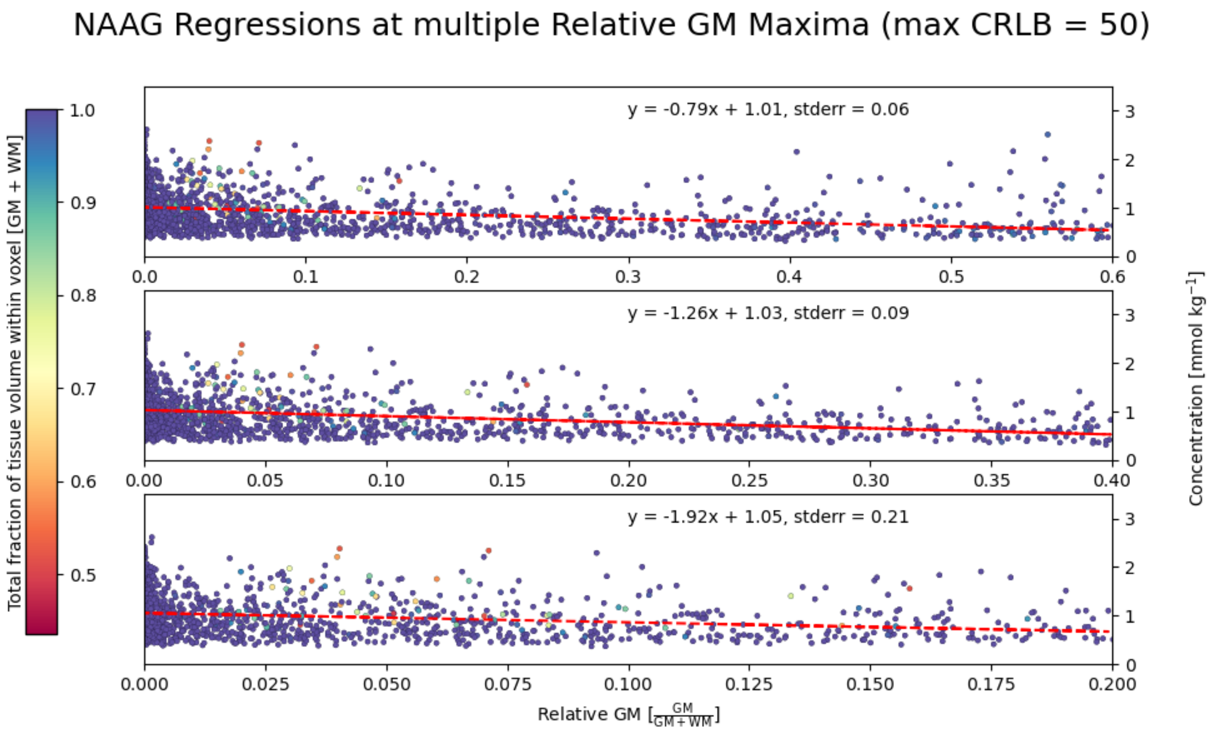

NAAG Regressions at multiple Relative GM Maxima (max CRLB = 100)

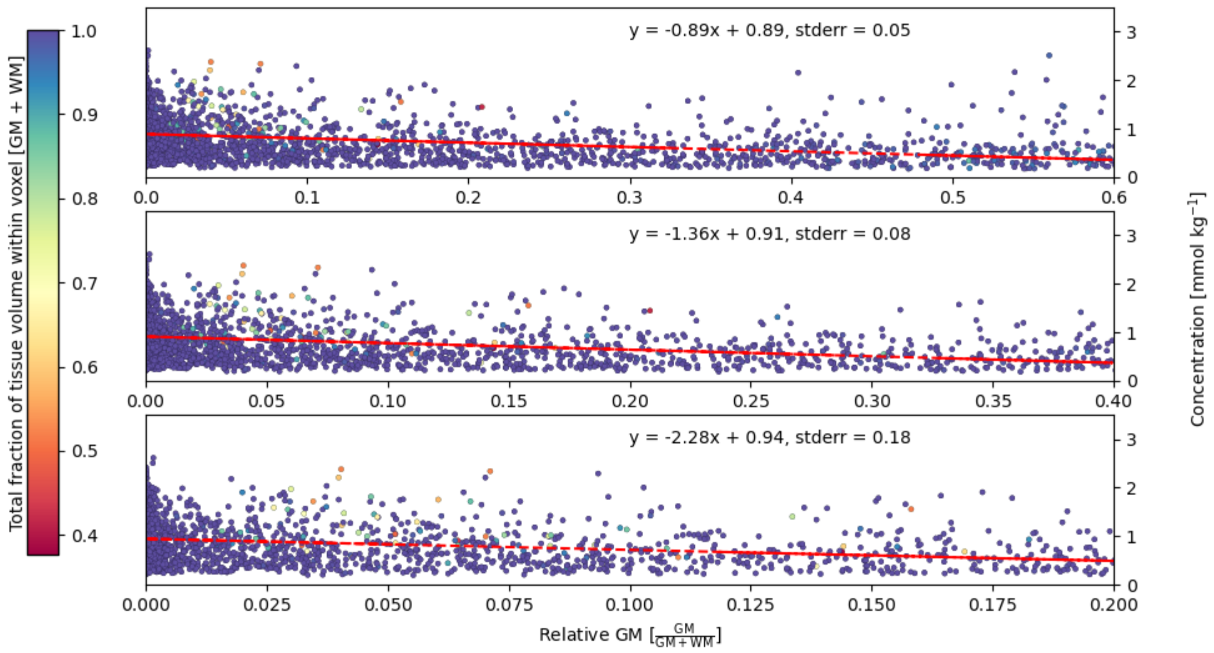


NAAG Regressions at multiple Relative GM Maxima (max CRLB = 500)

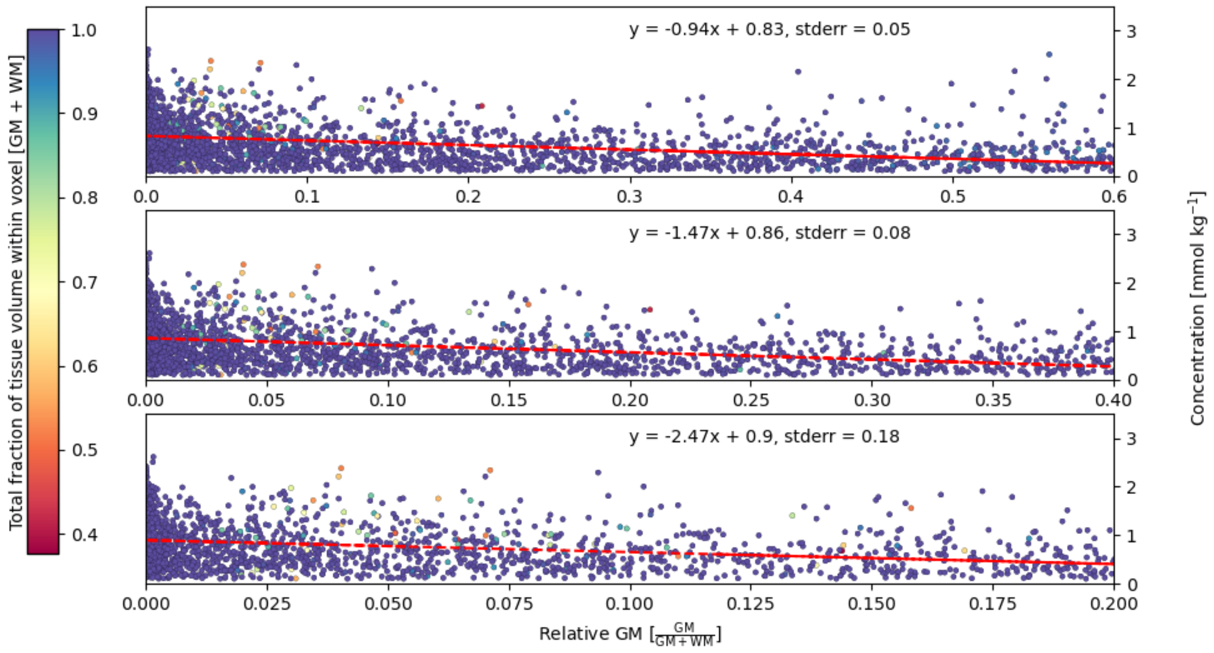

Supporting Information Table 1: continuation of Table 1 from the main text. Including which reference standard and relaxation corrections were used as well as how macromolecules were accounted for.

| Comparisons of 9.4 T $^{1}$H MRSI processing parameters to SVS studies [mmol kg$^{-1}$] | | | | | |
|---|---|---|---|---|---|
| | Region | $B_0$ (T) | Internal Reference | Relaxation corrections | Macromolecules |
| **This work** | GM-rich* | 9.4 | Water | voxel-specific $T_1$ for water and metabolites | $MM_{AXIOM}$ |
| **This work** | WM-rich* | 9.4 | Water | voxel-specific $T_1$ for water and metabolites | $MM_{AXIOM}$ |
| **Deelchand et al., 2010** | occipital lobe | 9.4 | Water | Not specified | measured |
| **Mangia et al., 2006** | occipital lobe | 7 | Assuming 80% water content | Not specified | measured |
| **Marjańska et al., 2012** | occipital lobe, mixed tissue | 7 | Water | $T_1$- and $T_2$-corrections for water. Only $T_2$-corrections for metabolites | measured |
| **Marjańska et al., 2012** | motor cortex, WM-rich | 7 | Water | $T_1$- and $T_2$-corrections for water. Only $T_2$-corrections for metabolites | measured |
| **Mekle et al., 2009** | occipital lobe, GM-rich | 7 | Assuming 80% water content and tCr = 8 mmol kg$^{-1}$ | Not specified | measured |
| **Murali-Manohar et al., 2020** | occipital lobe, GM-rich | 9.4 | Water | $T_1$- and $T_2$-corrections for water and metabolites | measured |
| **Terpstra et al., 2009** | occipital lobe | 7 | Assuming 80% water content | Not performed | measured |
| **Wright et al., 2021a** | occipital lobe, GM-rich | 9.4 | Water | $T_1$- and $T_2$-corrections for water and metabolites | $MM_{AXIOM}$ |
| **Wright et al., 2021a** | occipital lobe, WM-rich | 9.4 | Water | $T_1$- and $T_2$-corrections for water and metabolites | $MM_{AXIOM}$ |
| **Baker et al., 2008** | WM regions averaged | 3 | Water | $T_1$- and $T_2$-corrections for water and metabolites | No correction |
| **Baker et al., 2008** | GM regions averaged | 3 | Water | $T_1$- and $T_2$-corrections for water and metabolites | No correction |
| **Michaelis et al., 1993** | parietal lobe, GM-rich | 2 | Water but used external reference for quantification | $T_2$-corrections applied for metabolites | Not specified |
| **Michaelis et al., 1993** | parietal lobe, WM-rich | 2 | Water but used external reference for quantification | $T_2$-corrections applied for metabolites | Not specified |

**Annex A**: Fit settings used for fitting $^1$H FID MRSI data in LCModel. Each voxel was fitted individually, and thus, the suffix “row_0_col_0” would change for each voxel fitted.

```
$LCMODL
OWNER='Max Planck Institute Biological Cybernetics'
TITLE='MRSI quantitative data'
FILBAS=' ~/MRSI_quantitative_basis.basis'
FILRAW='~/met_RAW/subj_row_0_col_0.RAW'
FILH2O='~/wref_RAW/subj_row_0_col_0_wref.RAW'
FILPS='~/OUT/subj_row_0_col_0.ps'
FILCSV='~/OUT/subj_row_0_col_0.csv'
FILCOO='~/OUT/subj_row_0_col_0.coord'
LPS = 8
LCOORD = 9
LCSV = 11
ppmst= 4.2
ppmend= 1.8
nunfil= 2048
neach= 100
dows= T
dkntmn= 0.25
deltat= 1.25e-04
degzer= 0.0
degppm = 0.0
sddegz= 2.0
sddegp= 1.0
nnot2= 6
chnot2(1) = 'Lip13c'
chnot2(2) = 'Lip13d'
chnot2(3) = 'Lip13e'
chnot2(4) = 'Gua'
chnot2(5) = '-CrCH2'
chnot2(6) = 'MM20'
wconc= 55510.0
rfwhm= 1.45
hzpppm= 399.976
WSMET= 'tCr'
shifmx(2)= 0.01
shifmn(2)= -0.01
$END
```
